# Hilltopping influences spatial dynamics in a patchy population of tiger moths

**DOI:** 10.1101/2021.12.17.473236

**Authors:** Adam Pepi, Patrick Grof-Tisza, Marcel Holyoak, Richard Karban

## Abstract

Dispersal is a key driver of spatial population dynamics. Dispersal behavior may be shaped by many factors, such as mate-finding, the spatial distribution of resources, or wind and currents, yet most models of spatial dynamics assume random dispersal. We examined the spatial dynamics of a day-flying moth species (*Arctia virginalis*) that forms mating aggregations on hilltops (‘hilltopping’) based on long-term adult and larval population censuses. Using time-series models, we compared spatial population dynamics resulting from empirically-founded hilltop-based connectivity indices, and modeled the interactive effects of temperature, precipitation, and density dependence. Model comparisons supported hilltop-based connectivity metrics over random connectivity, suggesting an effect of hilltopping behavior on dynamics. We also found strong interactive effects of temperature and precipitation on dynamics. Simulations based on fitted time series models showed lower patch occupancy and regional synchrony, and higher colonization and extinction rates when hilltopping was included, with potential implications for the probability of persistence of the patch network. Overall, our results show the potential for dispersal behavior to have important effects on spatial population dynamics and persistence, and we advocate inclusion of such non-random dispersal in metapopulation models.

## Introduction

Dispersal plays a critical role in the dynamics of spatially structured populations [1,2]. It can maintain genetic diversity, rescue declining populations, and allow recolonization of those that are locally extinct [3–5]. For extinction-prone populations, network-wide extinction risk is spread through loosely linked patches by infrequent dispersal events [3,4]. Conversely, a patch-network tightly linked through dispersal is at greater risk of network-wide collapse due to synchrony of local populations. ‘Connectivity’ is the broadly defined metric (reviewed by [6]) used to quantify the strength of these linkages [7]. Because connectivity measurements are intuitive and linked to population persistence, they are regularly used in reserve design and conservation planning to mitigate the effects of global change [8].

Despite their wide-spread application, inclusion of connectivity metrics in patch-based models has historically provided poor predictions of patch occupancy [9]. This is in part due to the simplifying assumptions of such models. For example, many classic connectivity metrics assume random movement, where successful immigration into neighboring patches is based solely on interpatch distance and dispersal ability [10,11]. However, empirical investigations of dispersal have repeatedly demonstrated that the likelihood of an individual arriving at any particular patch is far from random [12]; resource availability, conspecific densities, and densities of natural enemies can affect emigration and immigration decisions [13,14]. Moreover, the transition through matrix habitat between suitable patches can be limiting, and is often influenced by landscape features. For example, topography can impede movement of some species [15] or attract individuals of other species that lek on hilltops, thereby creating barriers or corridors for dispersing individuals [16,17]. Because of the challenges of empirically measuring dispersal behavior though observation, telemetry, or mark-recapture, it is rarely done and even more seldomly incorporated into patch-based models.

The overarching aim of this study was to assess the implications of non-random dispersal for spatially-structured populations. To achieve this, we studied the spatial and temporal dynamics of a patchily structured, diurnal moth population (*Arctia virginalis*; Lepidoptera: Erebidae) that exhibits hilltopping behavior. Hilltopping is a common mate-locating strategy, where flying insects aggregate on sites of topographical prominence like hilltops and ridges to increase the likelihood of encountering conspecifics of the opposite sex [18,19]. Because hilltopping behavior constrains movement through hilltop mating aggregations, it is a useful means of studying the consequences of non-random dispersal [16,20–22]. In our previous work, a landscape connectivity index was developed that incorporated hilltopping behavior [23] based on a mark-recapture study by the same authors [24]. However, the usefulness (or validity) of this connectivity index relative to traditional metrics was not tested. In the present study, we specifically compared the effects of three different connectivity indices, two of which incorporated the hilltopping behavior of *A. virginalis*, using five additional years of data than our earlier work.

We focused specifically on the population-level consequences of hilltopping, by comparing the dynamical effects of three different connectivity indices, two of which incorporated the hilltopping behavior of *A. virginalis*. First, using survey data, we examined the relationships between adult moth and caterpillar abundances in larval patches and hilltops, which form the empirical foundation for behaviorally-based connectivity indices. We also used time series models of caterpillar data in survey patches to compare traditional and behaviorally-based connectivity indices as predictors of population dynamics. Previous work in this system found that precipitation had strong effects on dynamics [23]. In the present study, we expand on this by including another climatic variable, temperature, which was found to effect predator-prey dynamics of *A. virginalis* and its ant predator [25]. We also examined the interactive effects of temperature and precipitation on population dynamics, and the role of density-dependence caused by a recently discovered virus [26]. Lastly, we constructed simulation models based on parameterized time series models using three connectivity indices to compare the consequences of random vs. non-random dispersal on patch dynamics.

## Methods

### Caterpillar censuses

We estimated *A. virginalis* caterpillar abundance at 12 habitat patches at BMR (Figure 1) using 15 years of census data, from 2007-2021. Censuses were conducted in the last week of March each year, by counting the number of caterpillars observed on haphazardly selected *Lupinus arboreus* bushes of similar size at each patch (n = 10 bushes, 2007–2011; n = 15 bushes, 2011–2019; n = 20 bushes, 2020-2021; detailed methods in [27]. For analyses of the caterpillar censuses, we averaged over the number of bushes sampled and used the average density of caterpillars per bush at each site as our response variable.

**Figure 1.**
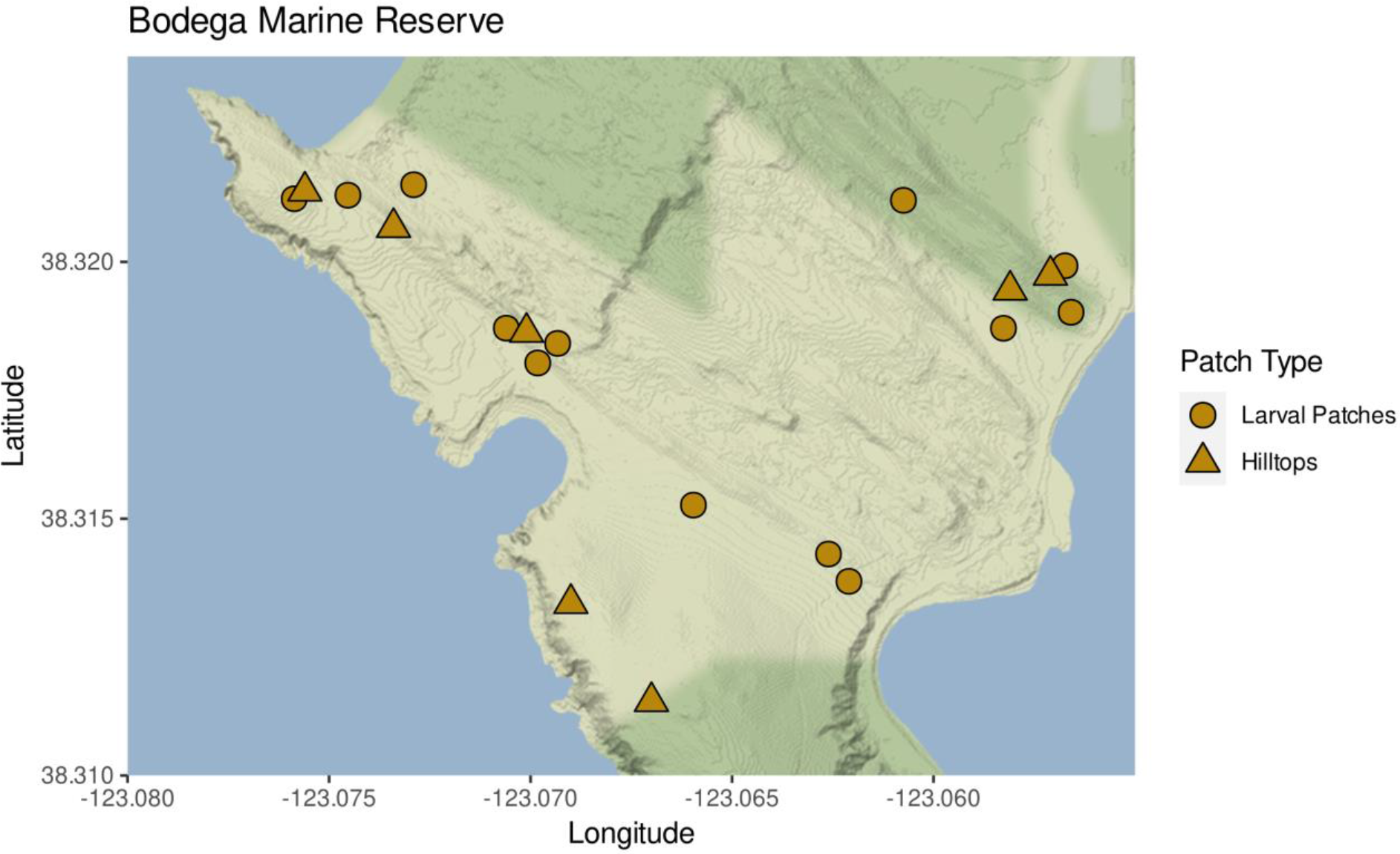
Map of Bodega Marine Reserve, showing location of surveyed larval patches (circles) and hilltops (triangles).

### Moth censuses

Moths at seven known hilltop lek locations [28] were surveyed annually from 2010-2012, and from 2017-2019. Surveys were not practical from 2013-2016 and 2020-2021 due to low adult abundances. Surveys were conducted at variable dates each year during the early summer, from 27 May - 19 July. This range of dates was required because the phenology of pupation varied among years. Surveys were timed roughly a month after pupation was observed. Adult moths are relatively long lived, persisting up to three weeks in mark-recapture and caging studies [24].

### Hilltops and larval patches

To investigate the determinants of hilltop aggregations, we analyzed predictors of moth abundance on hilltops based on prior work, including connectivity to larval patches, and elevation of hilltops [24]. In these models, we used we used a patch connectivity index similar to that used by [29], which was of the form:

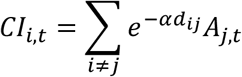

where *CI*_*i*_ is the connectivity index of patch *i* in year *t, α* is the inverse dispersal distance, which we set at 1 [23,24], *d*_*ij*_ is the distance between patch *i* and patch *j*, and *A*_*j*_ is log abundance in patch *j* in year *t*.

We constructed a set of models which we compared using WAIC, starting from the full model form:

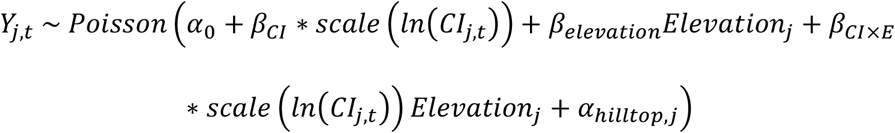

in which *Y*_*j,t*_ is the number of moths observed of hilltop *j* in year *t, CI*_*j,t*_ is a connectivity index of hilltop *j* in year *t* (logged and z-score scaled), *Elevation*_*j*_ is the elevation in ft of hilltop *j, β*_*CI*×*E*_ is an interaction between connectivity and elevation, and *α*_*hilltop,j*_ is a random intercept for hilltop. Models were fit using brms, using vague normal priors for all terms (*Normal*(0,10)), with *α*_*hilltop,j*_ having a hyperprior of mean *α* and variance *σ* both using the default brms priors, Student’s t (*t*(3,0,2.7)). Model structures without an interaction, with just elevation or connectivity, or an intercept-only model were compared with WAIC (Table 1).

**Table 1.** Results of moth abundance model structure comparison, showing Δ*WAIC* and *WAIC*.

| Model structure | $\Delta WAIC$ | $WAIC$ |
| --- | --- | --- |
| $X_t a_0 + \beta_{connectivity} + \beta_{elevation} + \alpha_{hilltop,j}$ | 0.00 | 296.33 |
| $X_t a_0 + \beta_{connectivity} + \alpha_{hilltop,j}$ | 6.58 | 302.91 |
| $X_t a_0 + \beta_{elevation} + \alpha_{hilltop,j}$ | 20.03 | 316.36 |
| $X_t a_0 + \alpha_{hilltop,j}$ | 22.56 | 318.89 |
| $X_t a_0 + \beta_{connectivity} + \beta_{elevation} + \beta_{CI \times E} + \alpha_{hilltop,j}$ | $4.3 \times 10^{37}$ | $4.3 \times 10^{37}$ |

To examine the influence of adult dispersal from hilltops on larval populations, we analyzed the relationship between moth abundance on hilltops and caterpillar abundance in larval patches in the subsequent year. We constructed a model of the following form:

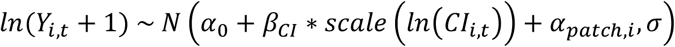

in which *Y*_*i,t*_ is the average caterpillar count in patch *i* in year *t, CI*_*i,t*−1_ is a connectivity index of patch *i* relative to hilltops in year *t-1* (logged and z-score scaled), and *α*_*patch,i*_ is a random intercept for patch. Models were fit using brms, using vague normal priors for all terms (*Normal*(0,10)) except the variance and random effects, with *α*_*hilltop,j*_ having a hyperprior of mean *α* and variance *σ*. Variance and hyperpriors were both the default brms priors, Student’s t (*t*(3,0,2.5)). The full model was compared to an intercept-only model using WAIC.

### Connectivity indices

For our analyses of caterpillar and moth abundances, we used several connectivity indices informed by *a priori* knowledge of the non-random dispersal behavior of adult moths. For a connectivity index representing random movement, we used the modified Hanski connectivity index defined above as *CI*.

We developed a connectivity index representing non-random movement occurring through hilltops, to test the role of hilltopping in driving dynamics. Our custom hilltop-based connectivity index was of the form:

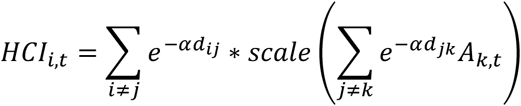

where *HCI*_*i,t*_ is the connectivity index of patch *i* in year *t, α*is the inverse dispersal distance, *d*_*ij*_ is the distance between patch *i* and hilltop *j, d*_*jk*_ is the distance between hilltop *j* and patch *k*, and *A*_*k,t*_ is log caterpillar abundance in patch *k* in year *t*. This connectivity index measures connectivity of patches through hilltops and was thus hypothesized to be a better representation of the spatial dispersal pattern of *A. virginalis*.

We also developed a connectivity index representing non-random movement occurring through hilltops that incorporated hilltop elevation. This was parameterized using the best fit model of moth abundance on hilltops (previous section), and was of the following form:

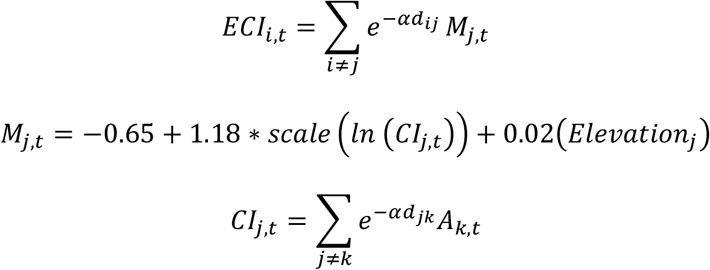

where *ECI*_*i,t*_ is the connectivity index of patch *i* in year *t, α*is the inverse average dispersal distance, *d*_*ij*_ is the distance between patch *i* and hilltop *j, M*_*j,t*_ is log predicted moth count on hilltop *j* in year *t, CI*_*j,t*_ is the connectivity index of hilltop *j* in year *t, Elevation*_*j*_ is the elevation in meters of hilltop *j, d*_*jk*_ is the distance between hilltop *j* and patch *k*, and *A*_*k*.,*t*_ is log caterpillar abundance in patch *k* in year *t*. This connectivity index measures connectivity of patches through hilltops, but with greater dispersal through higher hilltops, and was thus hypothesized to be a better representation of the spatial dispersal pattern of *A. virginalis* based on empirical findings from models of adult moth surveys (above sections).

### Caterpillar time series models

To examine the combined effects of density dependence, climate, and dispersal on caterpillar population dynamics, we constructed a series of time series models. We modelled dynamics in each patch separately but linked patches through dispersal via a connectivity term. We also included direct and delayed density dependence terms, intended to represent direct and delayed effects of baculovirus infection on dynamics that we have previously documented [26,30]. The model comparison started from the following full model structure:

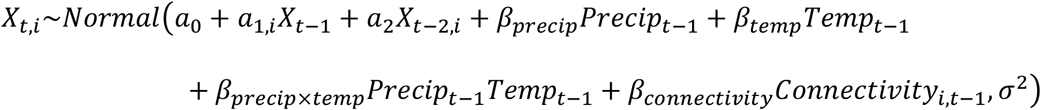

where *X*_*t,i*_ is the log caterpillar count in year *t* and patch *i*. The model includes an intercept representing population growth at low density (*a*_0_), direct density-dependence (*a*_1_), delayed density dependence (*a*_2_), an effect of precipitation (*β*_*precip*_), an effect of temperature(*β*_*temp*_), an interaction effect between precipitation and temperature (*β*_*precip*×*temp*_), and an effect of connectivity (*β*_*connectivity*_).

A model selection comparing models including all or a subset of parameters in the full model above was conducted (Table 2). Versions of all models using three different connectivity indices were compared: the standard patch connectivity index (*CI*), the hilltop-based connectivity matrix (*HCI*), or the hilltop-based connectivity index incorporating hilltop elevation (*ECI*; Table 2). Models were compared using WAIC. A prior of *Normal*(0,10) was used for all parameters except for the variance, for which we used a *Uniform*(0,10) prior. Models were fit in Stan [31].

**Table 2.** Results of autoregressive model selection, showing Δ*WAIC* from best model fit, *WAIC* values, Δ*WAIC* of the Hanksi connectivity index (*CI*) vs. the hilltopping connectivity index(*HCI*), and Δ*WAIC* of the elevation connectivity index (*ECI*) vs. the hilltopping connectivity index (*HCI*).

| Model structure | $\Delta WAIC$ | $WAIC$ | $CI - HCI$ | $\frac{ECI}{- HCI}$ |
| --- | --- | --- | --- | --- |
| $X_t \sim a_0 + a_1 + a_2 + \beta_{precip} + \beta_{temp} + \beta_{precip \times temp} + \beta_{connectivity}$ | 0.00 | 625.98 | 10.34 | 9.18 |
| $X_t \sim a_0 + a_1 + a_2 + \beta_{precip} + \beta_{temp} + \beta_{connectivity}$ | 17.05 | 643.03 | 5.46 | 8.49 |
| $X_t \sim a_0 + a_1 + a_2 + \beta_{temp} + \beta_{connectivity}$ | 20.47 | 646.45 | 8.70 | 9.09 |
| $X_t \sim a_0 + a_1 + a_2 + \beta_{precip} + \beta_{temp}$ | 25.97 | 651.95 | - | - |
| $X_t \sim a_0 + a_1 + a_2 + \beta_{temp}$ | 28.58 | 654.56 | - | - |
| $X_t \sim a_0 + a_1 + a_2 + \beta_{precip} + \beta_{connectivity}$ | 35.98 | 661.96 | -5.67 | 7.35 |
| $X_t \sim a_0 + a_1 + \beta_{precip} + \beta_{temp} + \beta_{precip \times temp} + \beta_{connectivity}$ | 39.43 | 665.41 | 1.66 | 8.37 |
| $X_t \sim a_0 + a_1 + a_2 + \beta_{precip}$ | 42.87 | 668.85 | - | - |
| $X_t \sim a_0 + a_1 + a_2 + \beta_{connectivity}$ | 56.14 | 682.12 | -12.52 | 5.03 |

### Simulation model

To assess the effects of non-random dispersal on spatial dynamics in this population, we conducted simulations using the best-fit time series model (Table 2). To conduct simulations, we used observed starting values for each patch in 2007 and 2008, and drew random values for population sizes in each patch for the subsequent year from a Poisson distribution using a mean value calculated with the time-series model formula. Empirical values of temperature and precipitation were used, and connectivity indices were calculated based on population sizes at each patch in the current year and scaled using the mean and standard deviation of observed connectivity values. We conducted versions of the simulations using the standard patch connectivity index (*CI*), the hilltop connectivity index (*HCI*), and the elevation connectivity index (*ECI*). Ten thousand simulations were run including the 15-year period, and occupancy, extinction, colonization, and regional synchrony (using package ncf [32] were summarized and compared between the different dispersal processes.

## Results

### Hilltops and larval patches

The best-fit model for adult moths on hilltops included connectivity to larval patches and elevation (Table 1), both of which were good predictors of moth abundance (Figure 2).

**Figure 2.**
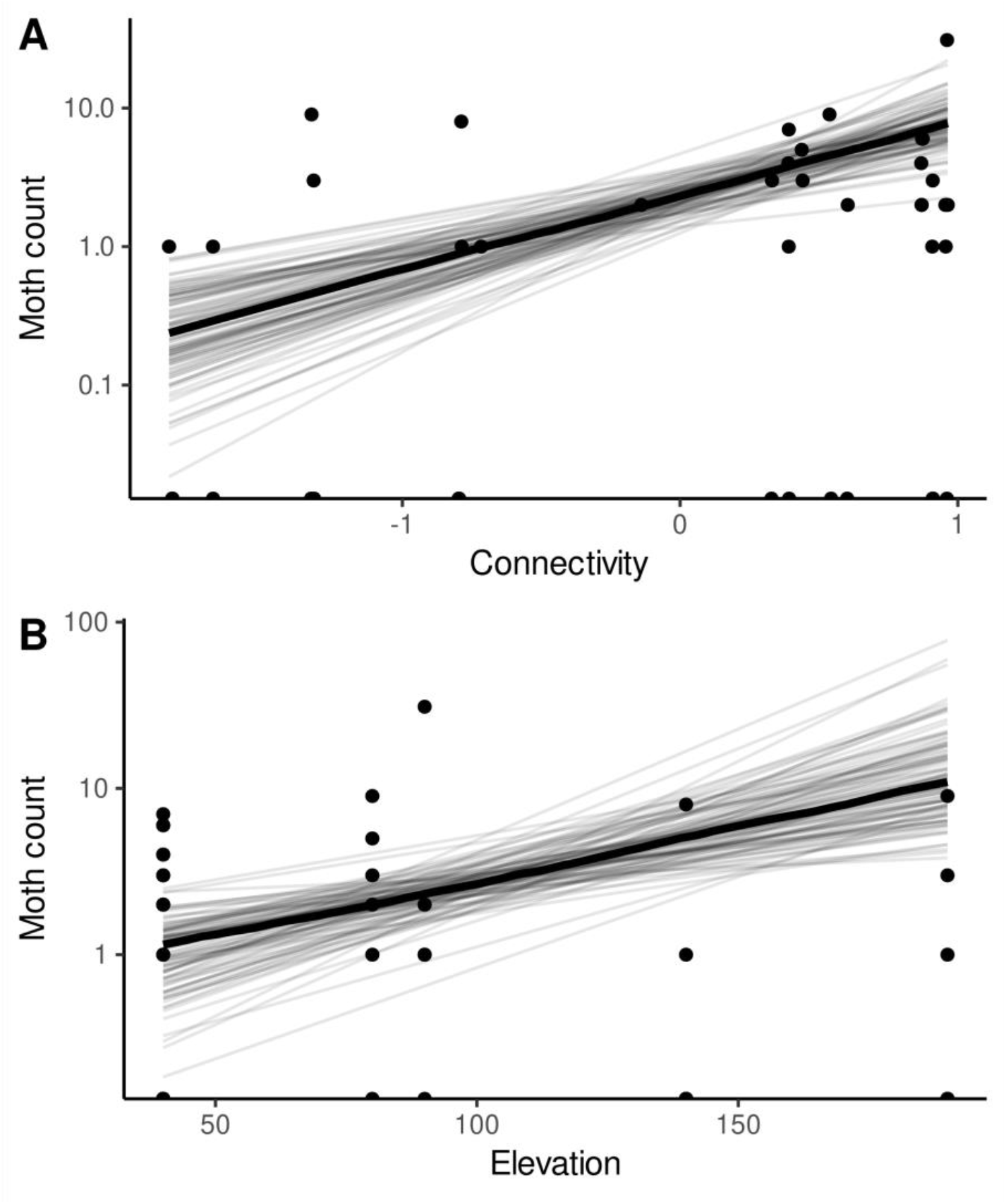
Best fit model of moth abundance, showing relationship with (logged and scaled) connectivity to larval patches (A) and elevation (B). Maximum *a posteriori* line is shown in wide black lines, plotted over 50 lines sampled from the posterior.

Connectivity between moths on hilltops and patches of larval habitat was a good predictor of caterpillar counts in patches in the subsequent year (*β*_*connectivity*_ = 0.5 [95%*HDPI*: 0.41 − 0.58], *R*^2^ = 0.14, Figure 3), and the model including connectivity had a better fit than the intercept-only model (Δ*WAIC*=1.8).

**Figure 3.**
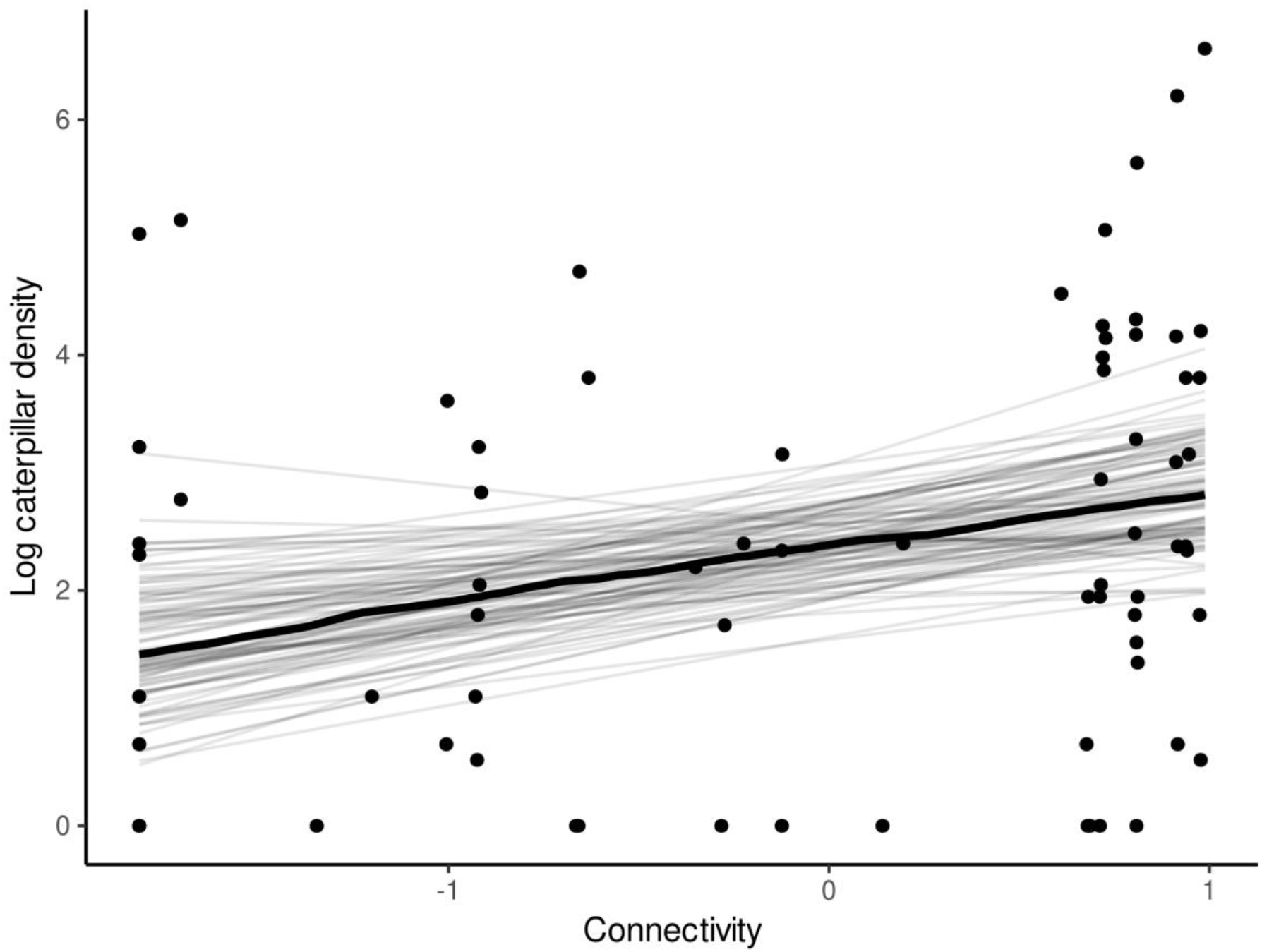
Model of caterpillar abundance, showing relationship with (logged and scaled) connectivity to moth hilltops. Maximum *a posteriori* line is shown in wide black lines, plotted over 50 lines sampled from the posterior.

### Caterpillar time series models

The best-fit model structure for the model comparison was the full model (Table 2), including direct and delayed density dependence, precipitation, temperature, connectivity, and an interaction between temperature and precipitation (Fig. 4). For the best-fit model, the hilltop connectivity index (*HCI*) was favored over Hanski’s connectivity index (*CI*) by Δ*WAIC*=10.34 and the elevation and hilltop connectivity index (*ECI*) by Δ*WAIC*=9.18 (Table 3).

**Figure 4.**
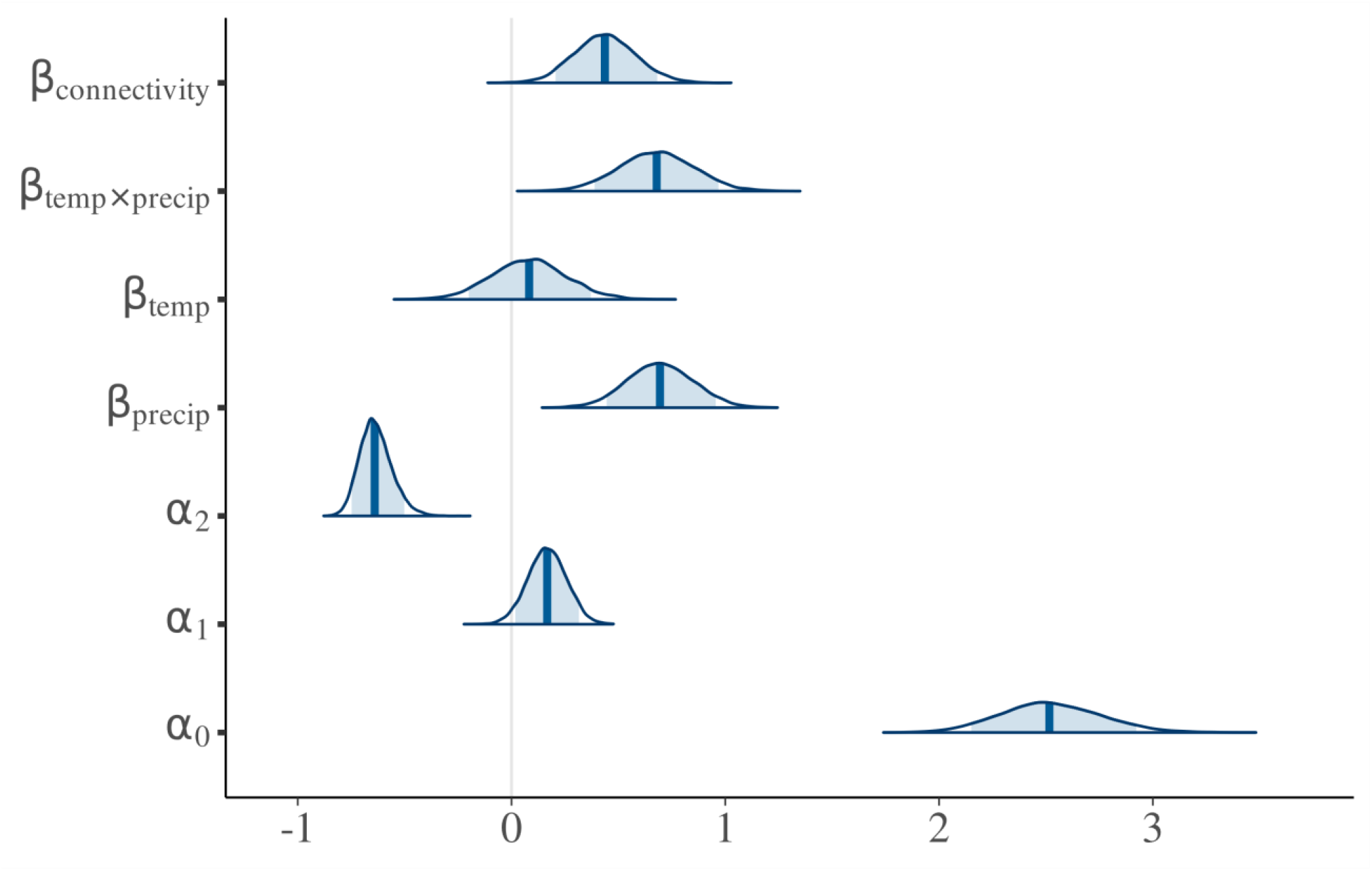
Posteriors densities from the best fit autoregressive model (table 2), with 90% intervals shaded.

### Simulation results

In metapopulation simulations, the hilltop connectivity index (*HCI*) and the elevation connectivity index (*ECI*) had somewhat higher colonization and extinction rates and lower occupancy compared to the standard patch connectivity index (*CI*) (Figure 5, Table S1), although simulated distributions overlapped substantially. The simulations with standard patch connectivity had higher regional synchrony than the hilltop or elevation connectivity models. For all metrics, hilltop-based connectivity simulation results had closer median values to observed values than other connectivity functions, although the actual observed values were quite different from the simulation results (Figure 5).

**Figure 5.**
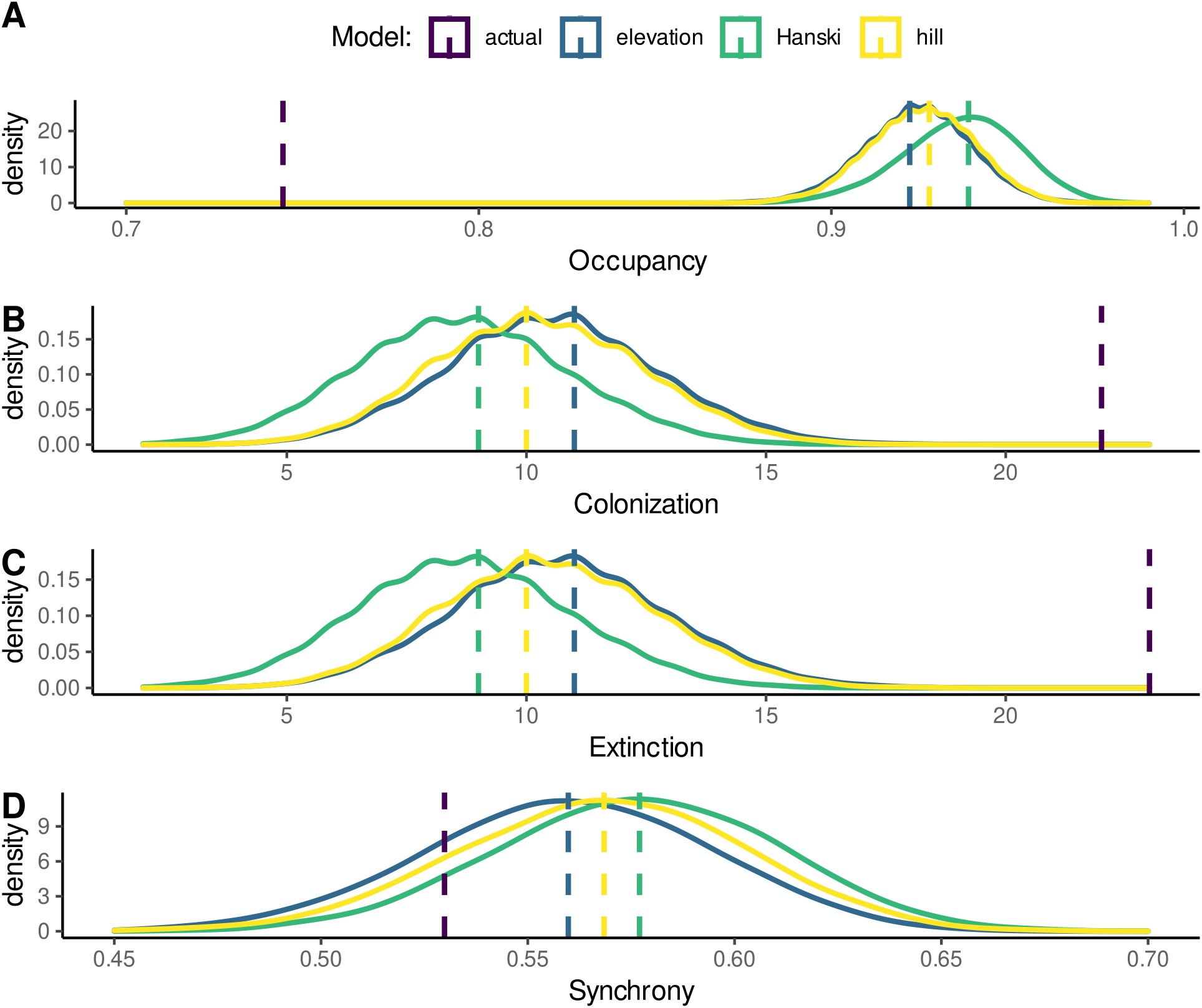
Connectivity simulation results, comparing occupancy (A), colonization (B), extinction (C), and regional synchrony (D) between the elevation, hilltop, and Hanski models.

## Discussion

Overall, our results showed that hilltopping behavior was a significant contributor to spatial dynamics in this population system. Empirically parameterized connectivity indices were significant predictors of population dynamics, and they were better predictors of dynamics than a standard connectivity index that did not include hilltopping (Table 2). Simulations demonstrated that, compared to other forms of connectivity function, hilltopping resulted in lower patch occupancy rates, higher extinction and colonization rates, and marginally lower regional synchrony (Fig. 5). Simulated outcomes from models including hilltopping were closer to observed results in the field than were results from simulations not including hilltopping. Our findings show that non-random dispersal behavior can influence spatial population dynamics, as has been predicted previously based on behavioral studies [12,16].

We also found empirical support for our hilltopping connectivity indices from monitoring moth abundances on hilltops. Results from models of hilltops and larval patches were consistent with observations that moths fly to nearby hilltops to find mates, and once mated, fly to nearby larval habitat patches and lay eggs. This local bi-directional movement produced a positive relationship between larval patches and nearby hilltop moth counts (Figure 2), and a similar positive relationship between hilltop moth counts and nearby larval counts in patches in the following year (Figure 3). Both estimates of larval and adult density are imperfect predictors of the next stage, since there is mortality (e.g., due to disease [30], predation [33]) during the intervening periods that is likely density-dependent. Such density-dependent processes have high potential to remove any correlation between these stages, and therefore the effects of movement were likely substantially stronger before density-dependent mortality. Overall, our analyses of these relationships gave strong empirical support for our hilltopping connectivity indices used in subsequent time series analyses.

Empirical comparisons of abundances on hilltops and larval patches found that connectivity indices that included hilltop elevation were better supported (Table 1), while time series models found more support for indices that included hilltopping but not elevation (Table 2). The latter result is also in contrast with previous findings that moth abundance on hilltops was well predicted by elevation [24]. A possible explanation for this discrepancy is that all of our empirical counts of adult moths have been conducted in relatively high-abundance years (2010-2012 and 2017-2019). Observations suggest that moths may be more likely to move on from local, lower elevation hilltops to taller more distant hilltops in higher density years (though we have surveys from too few years to test this formally). This may be due to high competition for mates at local hilltops near larval patches, which might be alleviated by dispersal in high density years (though the role of density in hilltopping behavior is unclear; see e.g., [18,34]). It is possible therefore that because of variation between years in the importance of elevation for aggregation size, the overall effect of elevation in both low- and high-density years was small enough such that it was not supported in our time series model selections.

Time series models suggested that the effects of connectivity were similar in magnitude or only slightly weaker than climate or density-dependent effects (Figure 4). Simulations further illustrated differences in dynamics between random movement, and dispersal that occurred specifically through hilltops. Because movement through hilltops is non-random and spatially restricted, simulations that included dispersal through hilltops found lower occupancy and higher extinction and colonization rates. This suggests that turnover (extinction and colonization) of local populations distant from hilltops may be more frequent than would be predicted by random movement models. In contrast, simulations with hilltopping dispersal had lower regional synchrony which could instead increase the likelihood of regional persistence [35]. This lower regional synchrony is likely due to hilltops serving as local attractors, driving divergence of dynamics between clusters of patches around different hilltops.

This study is the first empirical example demonstrating dynamical consequences of hilltopping in a patchy population, though previous work has shown effects of other modes of non-random dispersal on population dynamics [36]. Previous empirical and theoretical workers have hypothesized that non-random dispersal via hilltops will have strong effects on spatial population dynamics since these dynamics are heavily influenced by dispersal [16,21,22,37]. However, this question has not been tested directly: one study found that topographic prominence (i.e., hilltops) was a strong predictor of genetic differentiation in a hilltopping butterfly species (*Papilio machaon*), which would be an expected result of hilltops functioning as local attractors of population dynamics and dispersal [38].

We found strong interactive effects of temperature and precipitation on population dynamics, with higher population growth in wetter and warmer years (Fig S1). One potential explanation for this finding is that climate effects were mediated from the bottom up, via poor host-plant quality in years unsuitable for plant growth (hot and dry, cold and wet, etc.). Indeed, our current work suggests this may be the primary mechanism through which temperature and precipitation affect dynamics of early instar larvae in this population (Pepi, Grof-Tisza, Holyoak and Karban, unpublished data). This in agreement with other studies that inferred extinction and colonization dynamics were affected by climate-mediated changes in resource abundance or patch quality [23,39,40]. Considering that climate models predict that severe weather will become more frequent in California [41], finding that *A. virginalis* is sensitive to climatic variation has strong implications for the persistence of local populations; climate-mediated deterioration of habitat quality combined with fewer immigration events increases the likelihood of prolonged absences in isolated patches far from adult aggregations on hilltops. Though we demonstrated improved model performance when we used connectivity indices based on hilltoping behavior, temperature and precipitation were found to be stronger drivers than connectivity of the spatial and population dynamics of *A. virginalis*. Despite the dominance of the area-and-isolation paradigm, this work illustrates the importance of considering climate effects when modeling the dynamics of patchy populations [23,39,42].

Most if not all organisms disperse in some predictable way whether deliberately like migrating fowl or with less agency such as ballooning spiders adrift in the prevailing wind. Lekking mating systems, which have evolved in numerous taxonomic groups, are but one example of how a life-history trait may lead to non-random dispersal. Indeed, despite its widespread use, the assumption of random dispersal is rarely accurate. The advancement of landscape ecology has seen the development of better tools to predict movement patterns; however, many metapopulations models still rely on this outdated approach. Though our hilltopping connectivity index generated simulated results only modestly closer to observed results than a standard patch connectivity index, small differences may be important for species of conservation concern. Alarmingly, it is for these vulnerable species that metapopulations theory is often applied to identify critical habitat and to aid in reserve design. In these cases, empirical measures of dispersal behavior and patterns that increase realism along with realistic connectivity indices may be of utmost importance.

## Conclusion

Hilltopping is a common behavior for many butterfly species and other insects. In the present study, we provide the first empirical evidence of an effect of hilltopping on spatial population dynamics. We found evidence supporting the inclusion of empirically-based connectivity metrics that incorporated hilltopping behavior over those that did not. Moreover, population models including density dependence and the interaction of temperature and precipitation were favored over those that were less complex. In simulations of spatial dynamics based on model fits with different dispersal modes, we found that models with hilltopping were closer to observed patch dynamics, and had lower occupancy and regional synchrony, and higher extinction and colonization rates. This could affect the probability of regional persistence, depending on the balance of influence on extinction and colonization rates vs. regional synchrony. Overall, our study suggests the importance of incorporating ecological realism, in this case non-random dispersal, when modeling the dynamics of a spatially structured population in an increasingly fragmented world.

## Acknowledgements

This study was conducted at Bodega Marine Reserve, and was supported by NSF-LTREB-1456225. We would like to thank Jaqueline Sones (Bodega Marine Reserve) for facilitating our fieldwork.

## Author Contributions

AP and PGT conceived the study, all authors contributed to conceptual development, PGT, AP, and RK collected the data, AP developed the models and conducted the analyses, AP and PGT drafted the paper, and all authors contributed critically to revision. All authors approved submission of the final manuscript.

## Data Accessibility Statement

All data and code associated with this study are available on Zenodo at DOI:10.5281/zenodo.5773245.

## Supplement

**Table S1.** Results of metapopulation simulations measured in mean occupancy, colonization, extinction and synchrony from 10,000 runs using three different dispersal processes. Median, 25% and 75% quantiles of distributions are listed.

| Model | Measure | 25% Quantile | Median | 75% Quantile |
| --- | --- | --- | --- | --- |
| Hanski connectivity index (*CI*) | Occupancy | 0.928 | 0.939 | 0.95 |
|  | Colonizations | 7 | 9 | 10 |
|  | Extinctions | 7 | 9 | 10 |
|  | Synchrony | 0.554 | 0.578 | 0.601 |
| Hilltop connectivity index (*HCI*) | Occupancy | 0.917 | 0.927 | 0.933 |
|  | Colonizations | 9 | 10 | 12 |
|  | Extinctions | 9 | 10 | 12 |
|  | Synchrony | 0.545 | 0.568 | 0.59 |
| Elevation connectivity index (*ECI*) | Occupancy | 0.917 | 0.927 | 0.933 |
|  | Colonizations | 9 | 10 | 12 |
|  | Extinctions | 9 | 11 | 12 |
|  | Synchrony | 0.536 | 0.560 | 0.582 |

**Figure S1.**
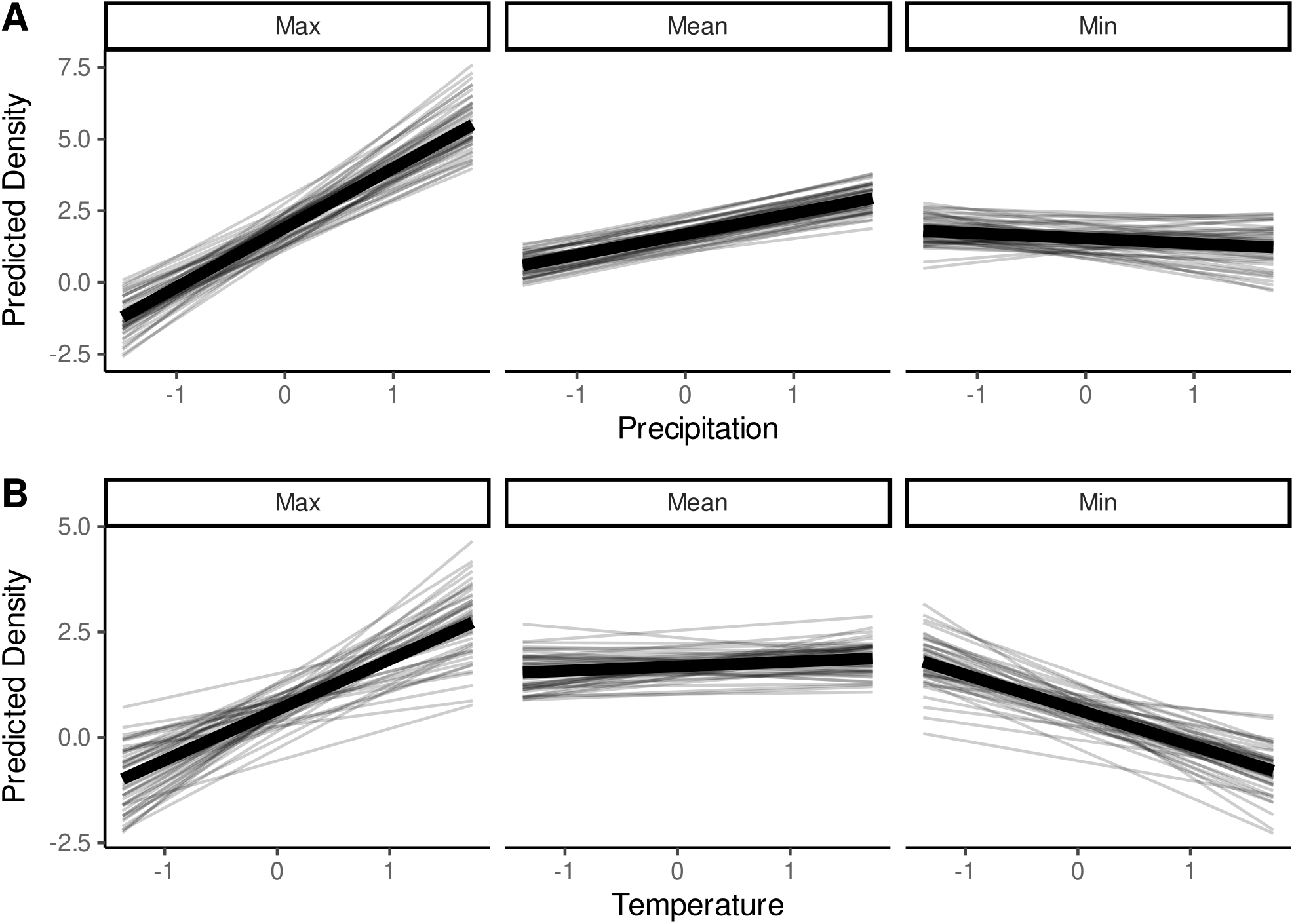
Plot of interaction effect between precipitation and temperature from best fit autoregressive model, showing predicted log population density depending on scaled precipitation (A) and temperature (B) at minimum, mean, and maximum values of temperature (A) and precipitation (B). Maximum *a posteriori* line is shown in wide black lines, plotted over 50 lines sampled from the posterior.

